## Supplementary figures and images for "Complementary CRISPR screen highlights the contrasting role of membrane-bound and soluble ICAM-1 in regulating antigen specific tumor cell killing by cytotoxic T cells"

### Figure 1

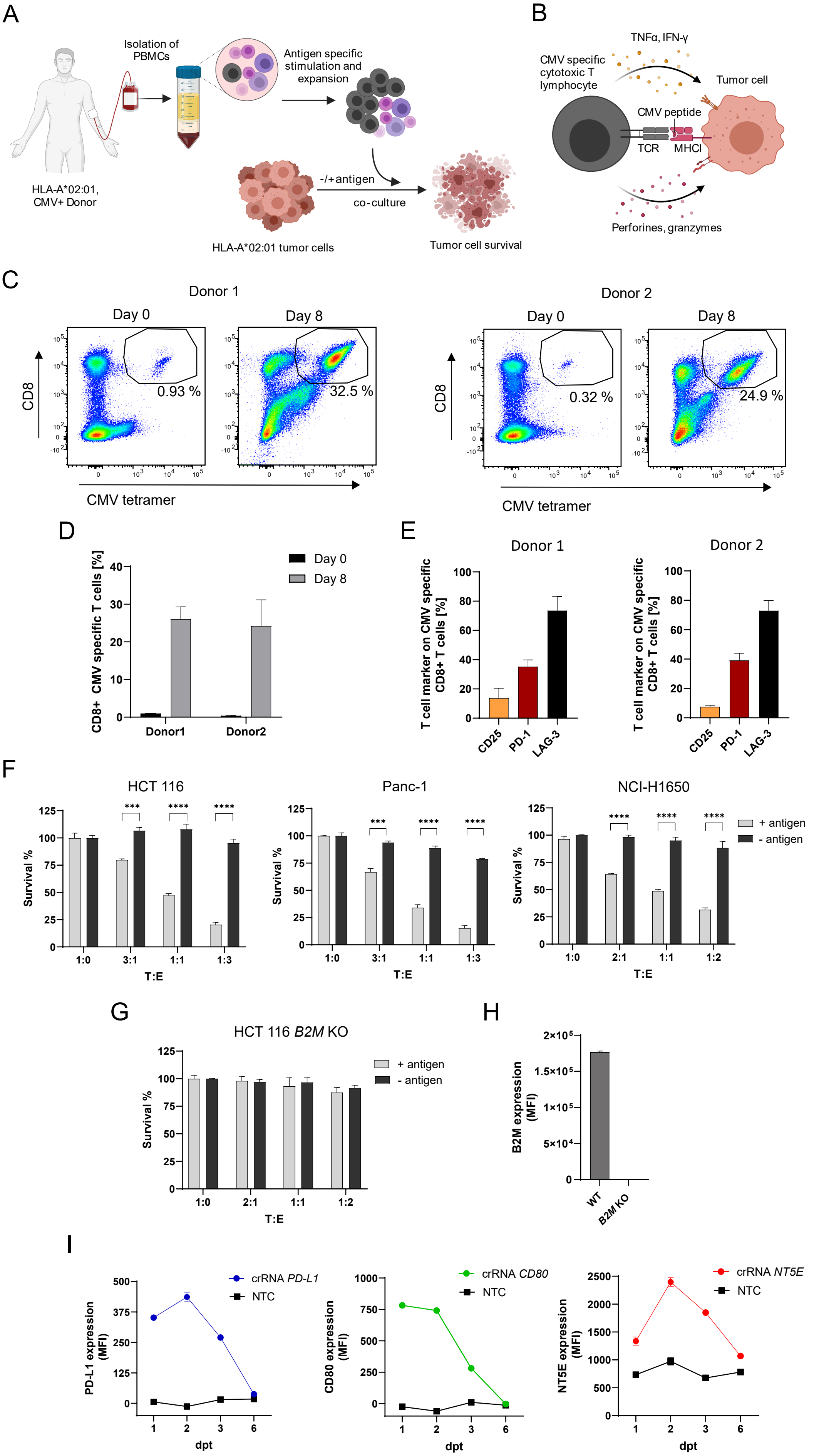

### Figure 2

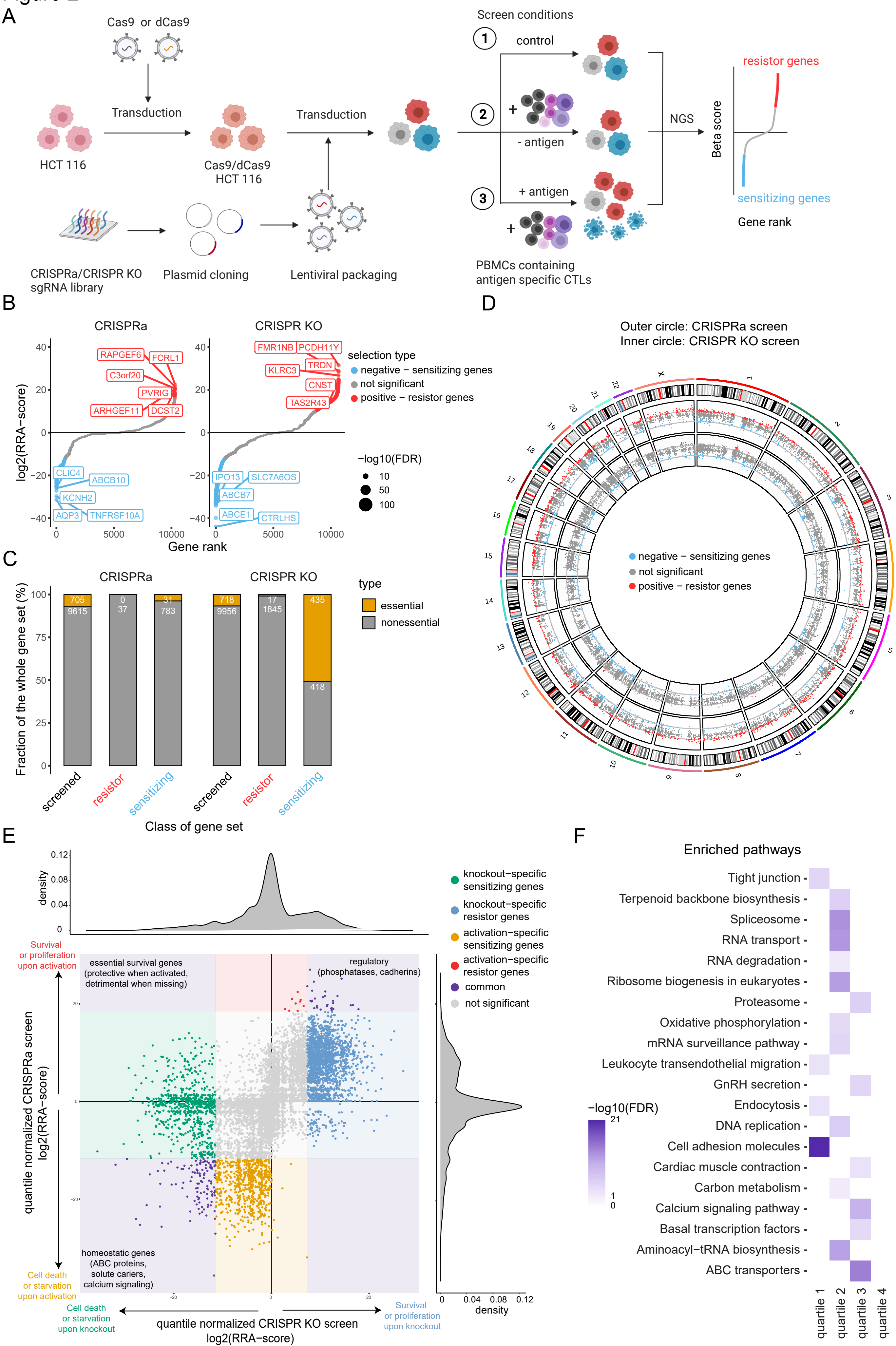

### Figure 3

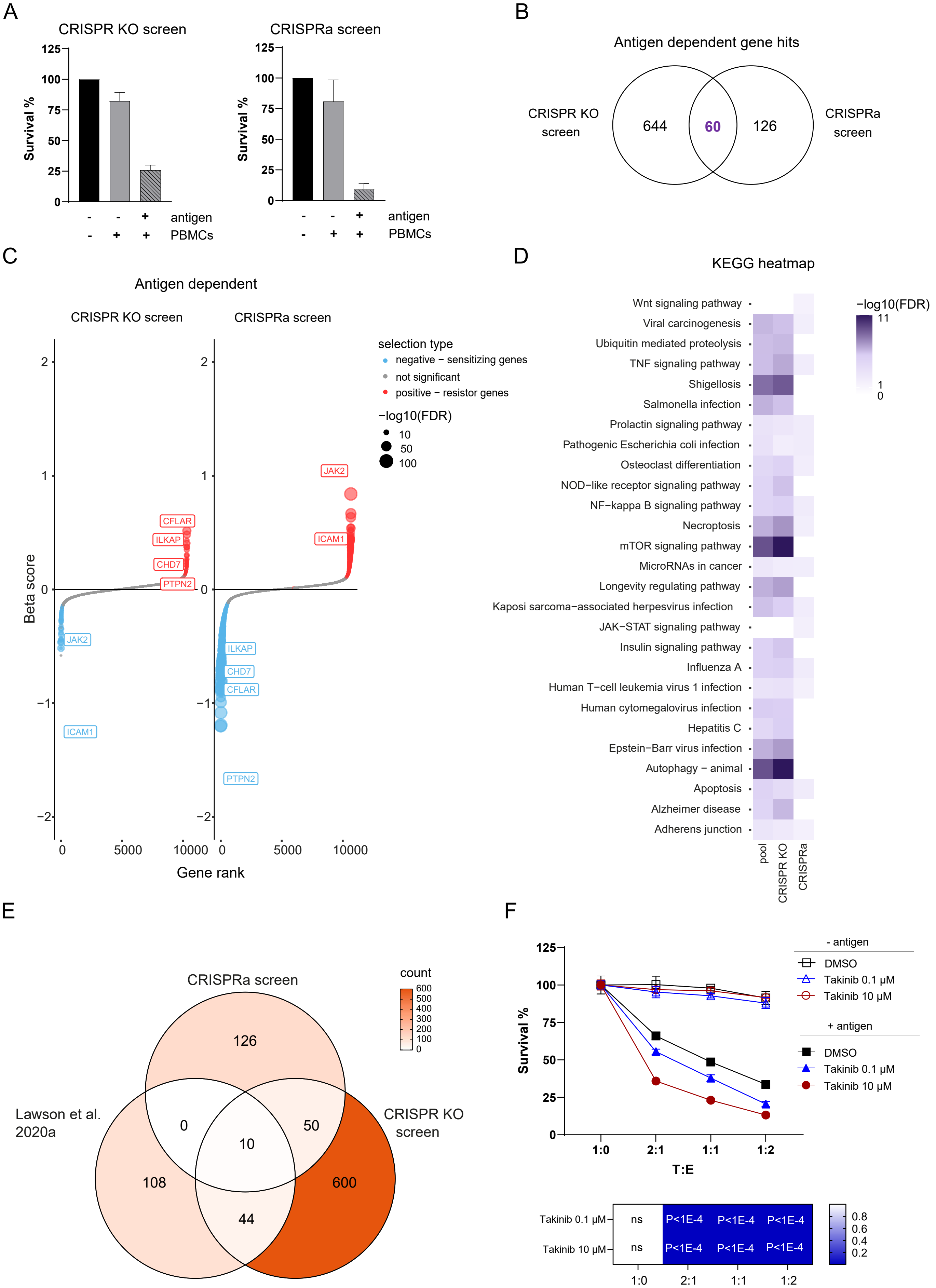

### Figure 4

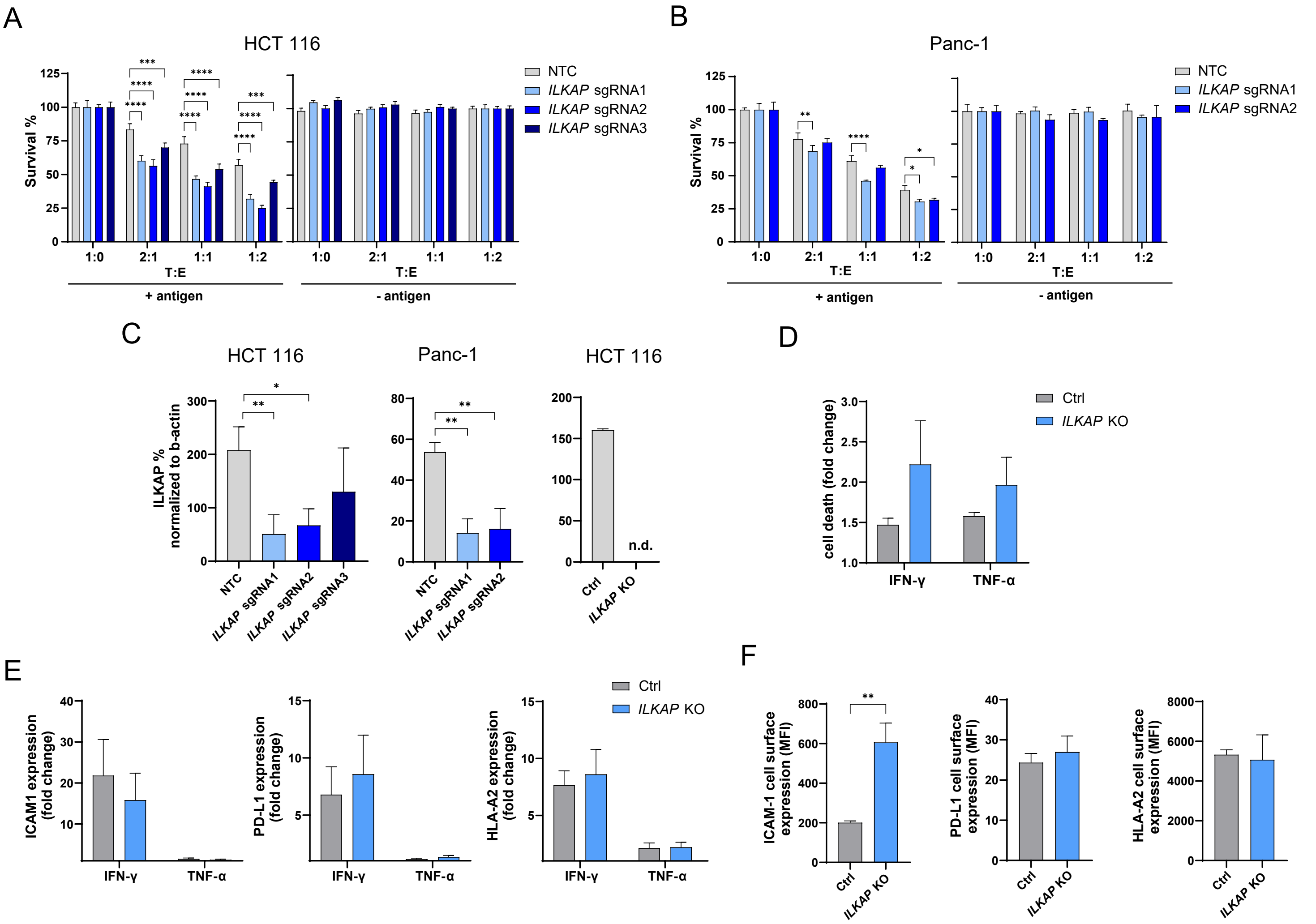

### Figure 5

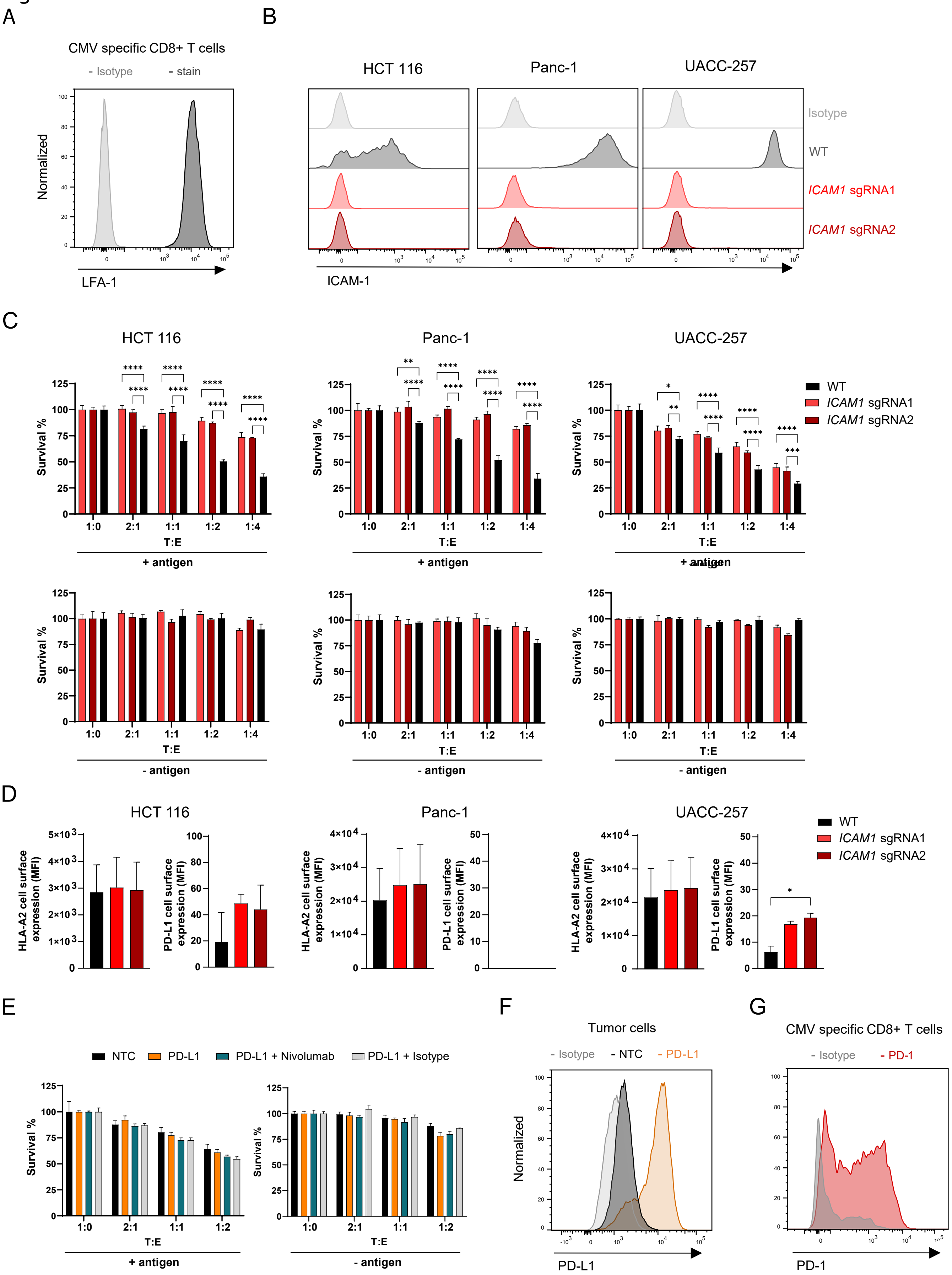

### Figure 6

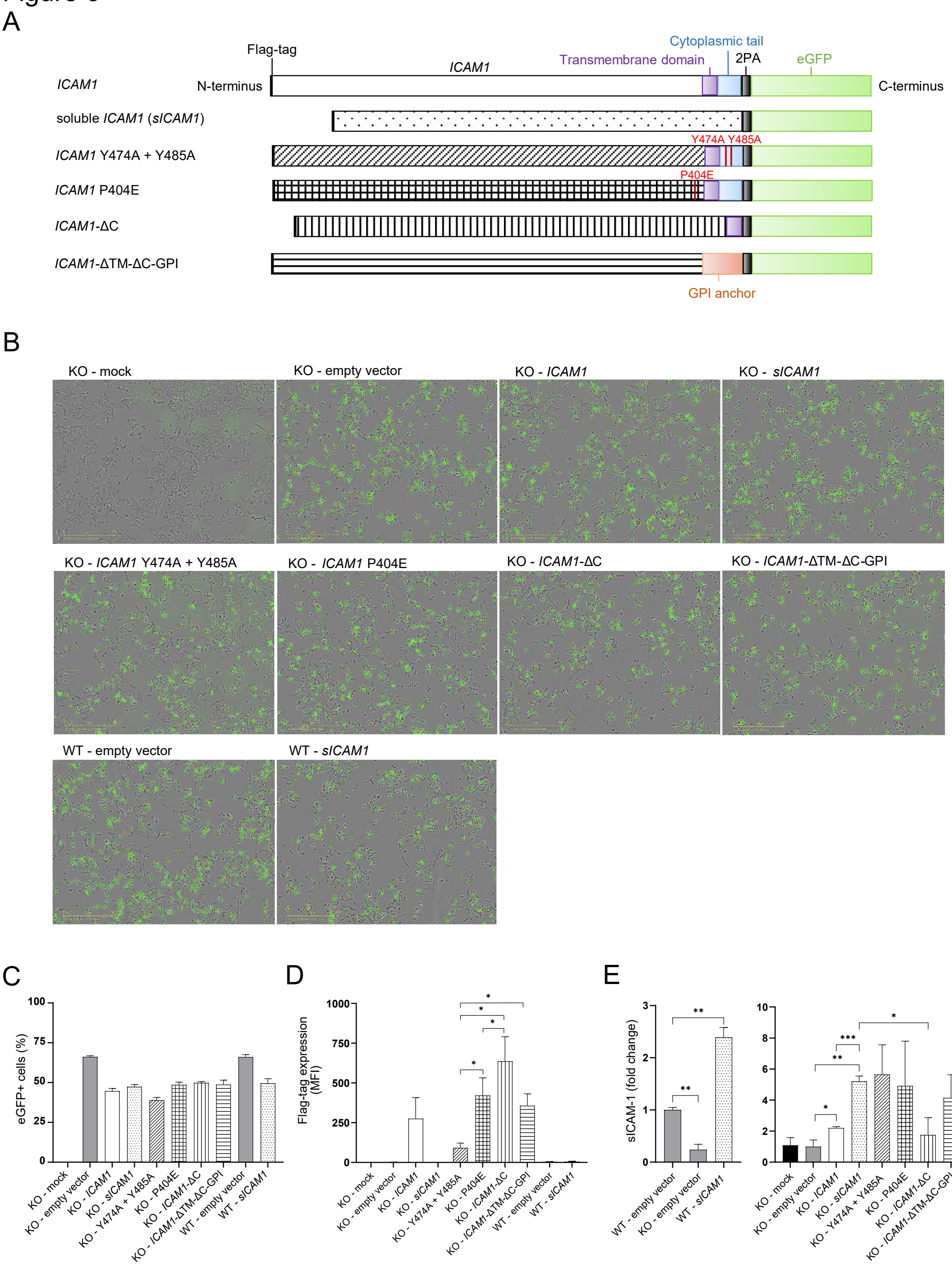

### Figure 7

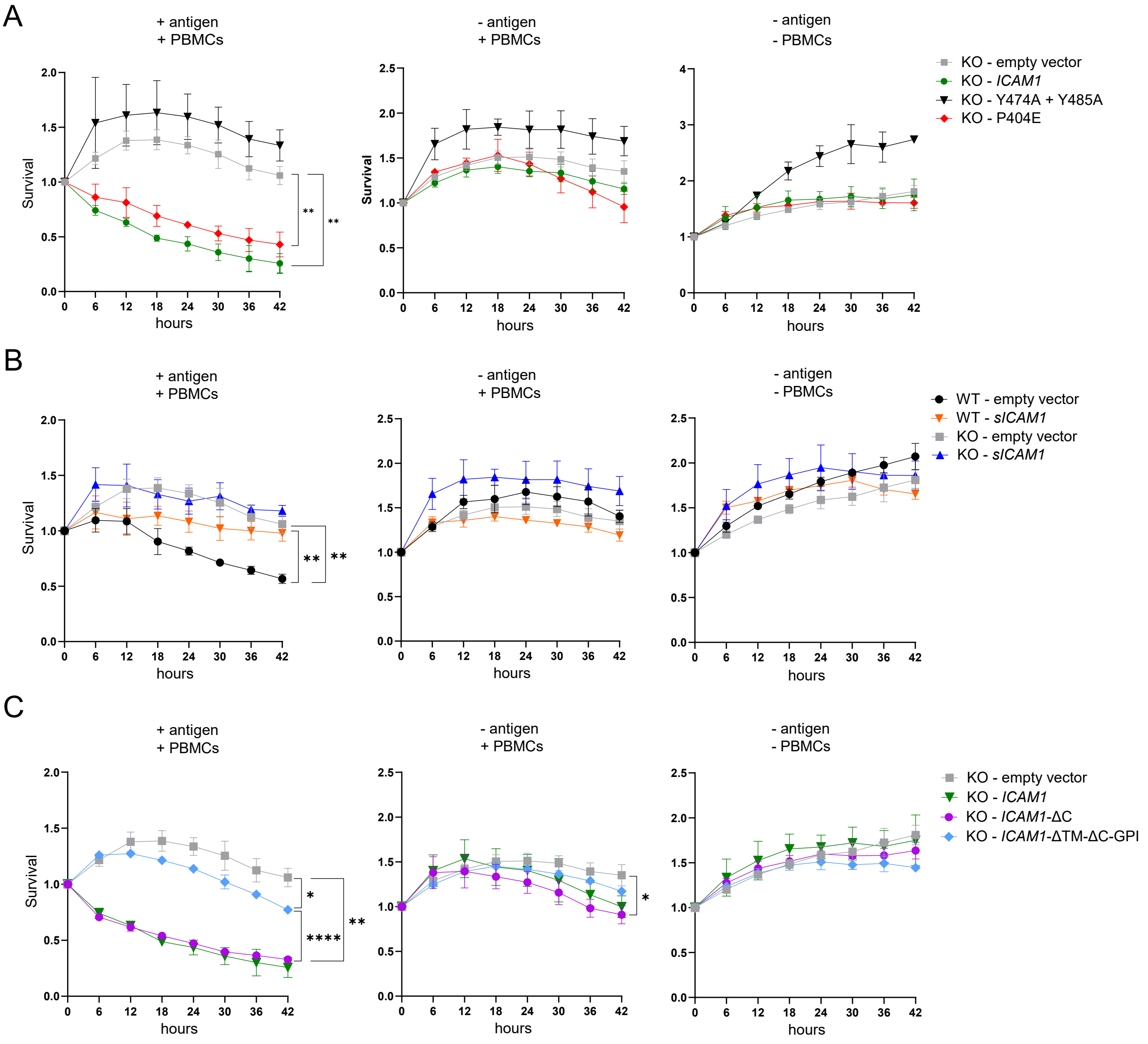

### Figure 8

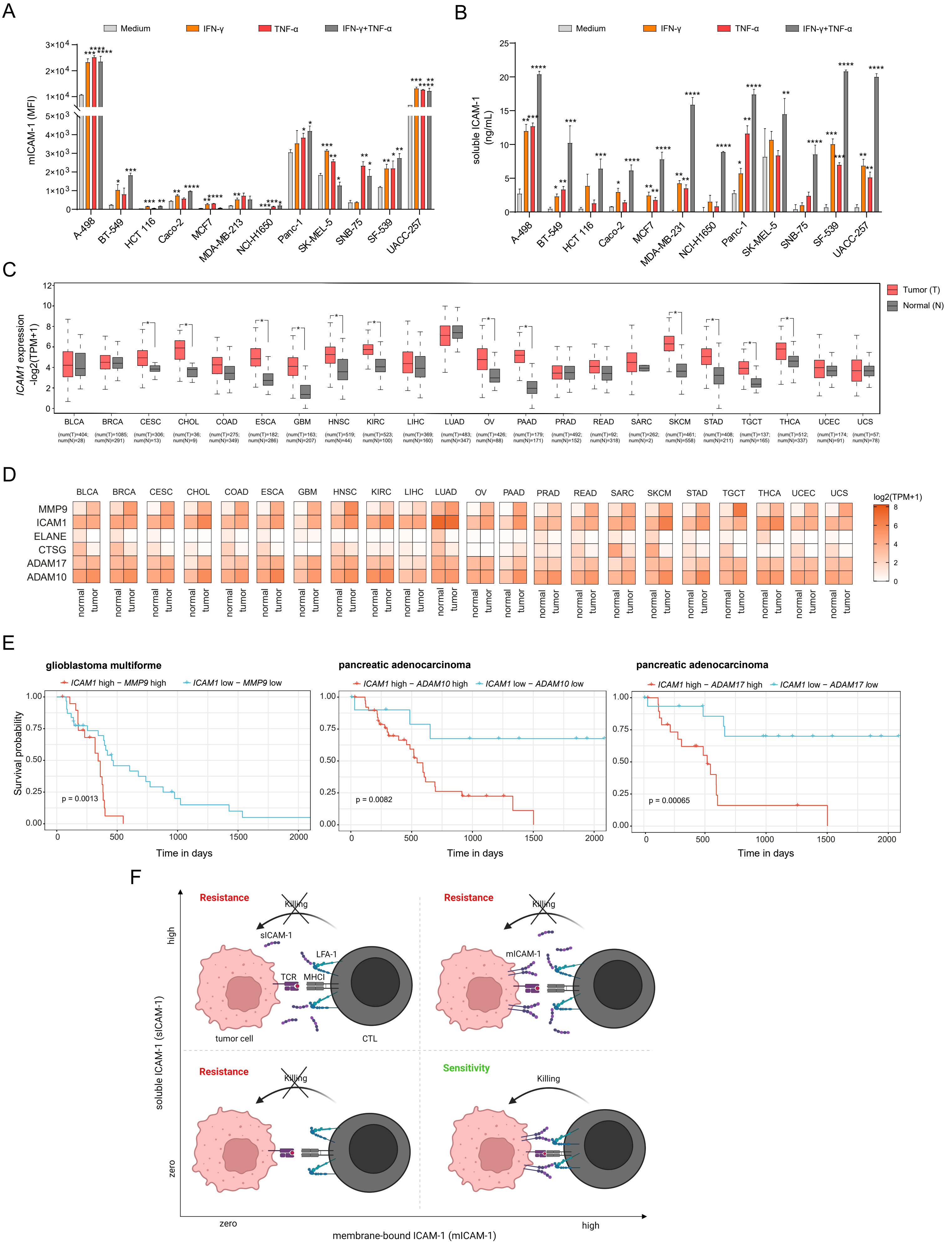

### Supplemental Figure 1

Suppl. Figure 1

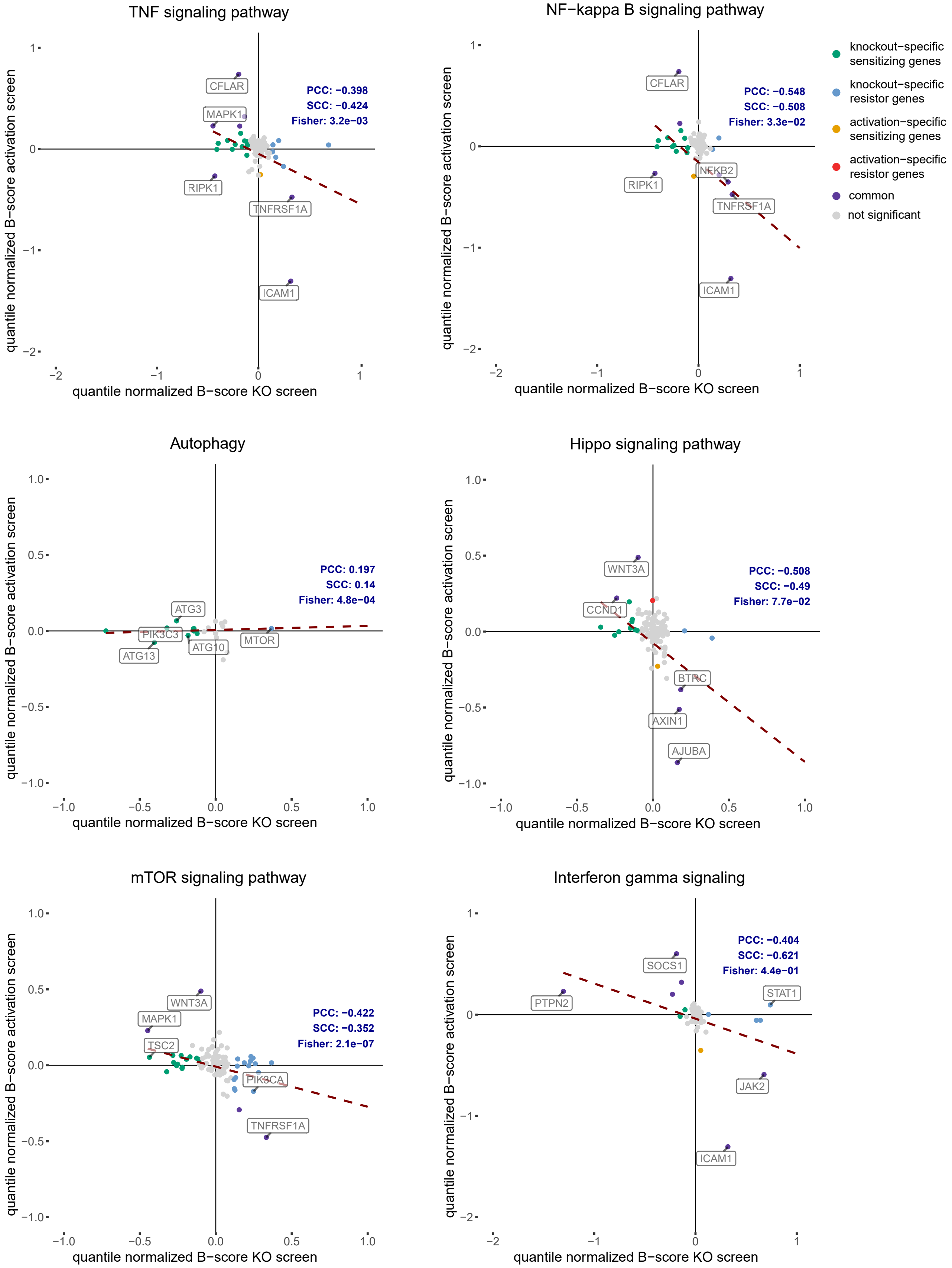
